# Photoelectricity generation from beet juice in a bio-electrochemical cell

**DOI:** 10.1101/2024.03.16.585328

**Authors:** Yair Farber, David Rasuli, Yaniv Shlosberg

## Abstract

The world is striving for the development of novel clean energy technologies that can replace the utilization of fossil fuels, and assist in the struggle against climate change. A unique approach that does not involve any carbon emission is bio-photo electrochemical cells (BPECs). This method integrates live organisms with an electrochemical system, while native redox species that exist in the organisms can generate dark and photo electrical current. Such systems were previously shown using photosynthetic micro and macro organisms in which the photocurrent mostly derives from photosynthesis. Another electron source was reported to originate from cherries which consist of pigments and enhance the production of the strong electron donor ascorbic acid under solar irradiation. In this work, we show the ability to produce electrical and photoelectrical current from beet juice. We apply cyclic voltammetry and fluorescence methods to show that among the major components that play a role in the current and photocurrent generation are NADPH and flavins.

## Introduction

In recent years, extensive efforts have been conducted to find alternatives for fossil fuels whose combustion increases the carbon emissions to the atmosphere. The idea of integrating a biological component within an electrochemical system was first applied in 1910 by Potter et al. who invented the first microbial fuel cell (MFC)[1]. Over the years, numerous improvements have been reported in the field of MFCs[2–4]. Among them is the utilization of bacterial species with a high exo-electrogenic activity such as *Shewanella Oneidensis*[5–7] and *Geobacter. Sp*.[8– 11] which can perform a direct electron transfer to the anode by *pili* or metal respiratory protein complexes. Bacterial cells can also generate electricity by a mediated electron transfer mechanism releasing redox species such as phenazines and quinones [12]. Interestingly, under irradiation, some of these molecules which are in their oxidized form may be photo-reduced enabling them to donate electrons at the anode of the bio-electrochemical cell (BEC)[13].

Another way to enhance the performance of BECs is the improvement of the electrochemical setup using advanced electrode materials with higher conductivity and Fermi energetic levels that are more compatible with the electron donors released by the cells[14–18]. The current generation may also be significantly enhanced by adding artificial redox species such as potassium ferricyanide which can mediate electrons from the external surface of the cells and the anode[19]. Rather than using bacteria, MFCs may also be based on yeast cells[20]. A major challenge for using MFCs is the maintenance of bacterial viability over time. To do that, nutrients and sugars should be constantly provided to feed the bacteria, which adds to the cost and complexity of the designed BEC. A way to overcome this obstacle is plant-MFCs in which the BEC is installed in the soil under plant roots[21,22]. The degradation of the roots causes the release of organic matter which can act as food for the bacteria keeping them alive. Furthermore, the roots also release redox species such as NADH and vitamins which can donate electrons at the anode regardless of the bacteria[23]. The maintenance of bacterial viability is much easier while working with photosynthetic bacteria or microalga cells, as they can synthesize their carbon source by photosynthesis[24]. Another advantage of using photosynthetic microorganisms is their ability to reduce carbon from the environment[25]. Upon illumination, photosynthetic microorganisms generate NADPH which can exist in the cells and donate electrons at the anode to enhance producing photocurrent[26–30]. Enhancement of the current in both photosynthetic and non-photosynthetic micro-organisms can be done by bio-film structures in which the cell density associated with the anode is bigger[10,31,32]. A photocurrent generation higher than micro-organisms can be produced by photosynthetic macro-organisms such as plant leaves[29,33] and seaweeds[34], in which the cells are organized in the form of a tissue that is denser than microbial suspensions. While non-biological electrochemical cells can tolerate extreme environmental conditions such as pH and salinity, BECs are more limited as the viability of the organism must be kept to enable a long-term current generation. A big advantage in BECs can be achieved while using marine organisms that can tolerate a high salinity which enables the use of a more conductive electrolyte[35]. Recent studies have shown the concept of using a fruit as an electron source producing dark and photocurrent from cherries and their juice[36]. In this work, we design a BEC based on beet juice, showing its ability to produce photocurrent in dark and under white light irradiation. Also, we apply cyclic voltammetry and fluorescence to reveal the identity of some of the major electron donors.

## Materials and methods

### Materials

All chemicals were purchased from Merck.

### Electrochemical measurements

All electrochemical measurements were conducted on a PalmSens4 potentiostat equipped with a commercially available screen-printed electrodes (SPEs) with graphite working and counter electrodes, and Ag coated with AgCl reference electrode. A drop of 50 µL of the analyte was placed on top of the SPE to cover the entire area of SPEs. Photocurrent measurements were done by placing a white light emitting diode (LED) above the SPEs. The light intensity at the SPEs surface height was measured by a light meter and set to 1000W/m2 by adjusting the distance of the light source from the SPEs. Chronoamperometry was measured applying an electrical potential of 0.9 V on the working electrode. Cyclic Voltammetry was conducted sweeping the potential from 0 to 1.2 V with a scan rate of 0.1 V/s.

### Fluorescence measurements

Prior to the measurement, the beet juice was diluted by a factor of 100,000 times. was conducted using a Cary Eclipse fluorimeter (Varian) with excitation and emission slits bands of 5 nm, applying of a voltage of 700 V on the photomultiplier detector.

## Results

### Studying the electrochemical activity of beet juice

Previous studies showed that cherry juice consists of various redox species that can donate electrons in the dark or act as photoelectron donors under white light illumination[36]. We wished to study the electrochemical activity of beet juice. To do this, we have designed a photo BEC system that consists of screen-printed electrodes (SPEs) with graphite working and counter electrodes, and silver coated with silver chloride reference electrode (Fig. 1a, b). A drop of 50 µL beet juice was squeezed directly from a live beet and placed on the SPEs. Light irradiation (1000 W/m^2^) was conducted from above using a white LED. Cyclic voltammetry was measured at the potential range of 0 – 1.2 V. The CV of the beet juice showed two major peaks around 0.7 V which is compatible with the voltammogramic fingerprint of anthocyanins and ascorbic acid[37], and around 0.9 V which may indicate the presence of the electron donor NADH[38]. These redox species were also previously reported to generate dark and photocurrent in cherries-based BECs. To eliminate the possibility the peaks originate from impurities of the system, CV of NaCl was measured showing no peaks.

**Fig.1.**
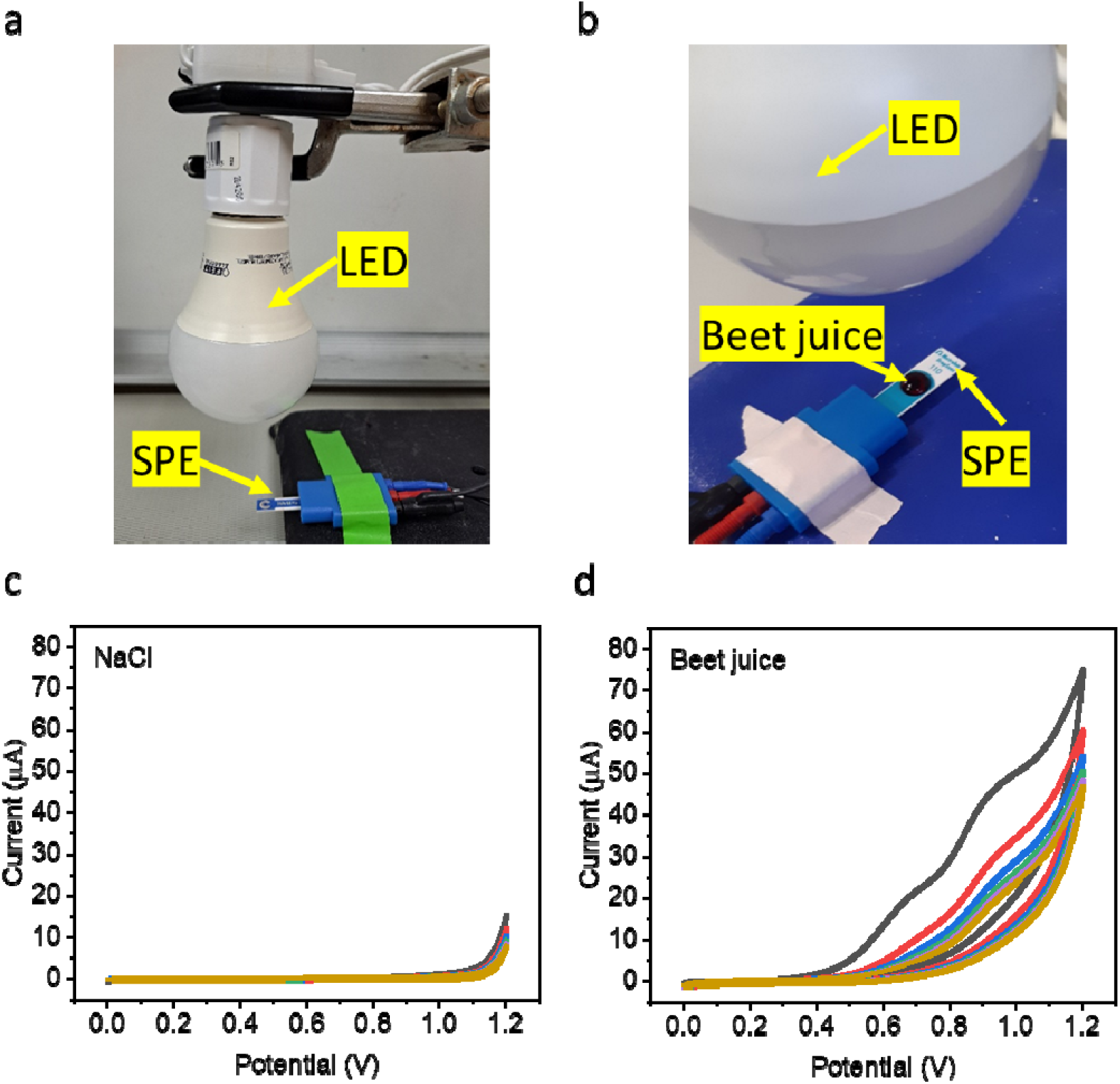
Studying the electrochemical activity of beet juice. CV of NaCl (control) and beet juice were measured (0 – 1.2 V) to detect redox-active species. **a** A picture of the measurement system that consists of a white LED above screen-printed electrodes (SPE). **b** A magnification of the system’s picture showing the beet juice drop placed on the SPE. **c** CV of NaCl solution. **d** CV of beet juice. Black, red, blue, azure and orange lines represent cycles 1-5 respectively.

### Spectral identification of fluorescent redox-active species

Previous BECs-related studies done on diverse cells[26,39–41] and organisms[33,34,42] such as cyanobacteria, microalgae, mammalian cells, seaweeds, plant leaves, and fruits have all showed reported the potential activity of redox-proteins, NADH, and flavines as electron donors in BECs. We wished to study if the same components also exist in beet juice. To elucidate this, fluorescence measurements were conducted exciting at wavelengths of 270, 350, and 450 nm which is compatible with the maxima of the amino acids tyrosine, tryptophan[43], NADH[26], and flavins[44] (Fig. 2a-c). The obtained spectra showed the spectral fingerprint of tyrosine and tryptophane, NADH, and flavins at maximal emission wavelengths of ∼ 330, 440, and 520 nm. Based on these peaks, we suggest that proteins, NADH, and flavins exist in the beet juice and may contribute to the current production of a BEC. The spectra excited at 350 nm have also shown additional peaks with a maximal emission at 500 and 600 nm. We suggest that these peaks may originate from the pigments in the beet juice such as anthocyanin.

**Fig.2.**
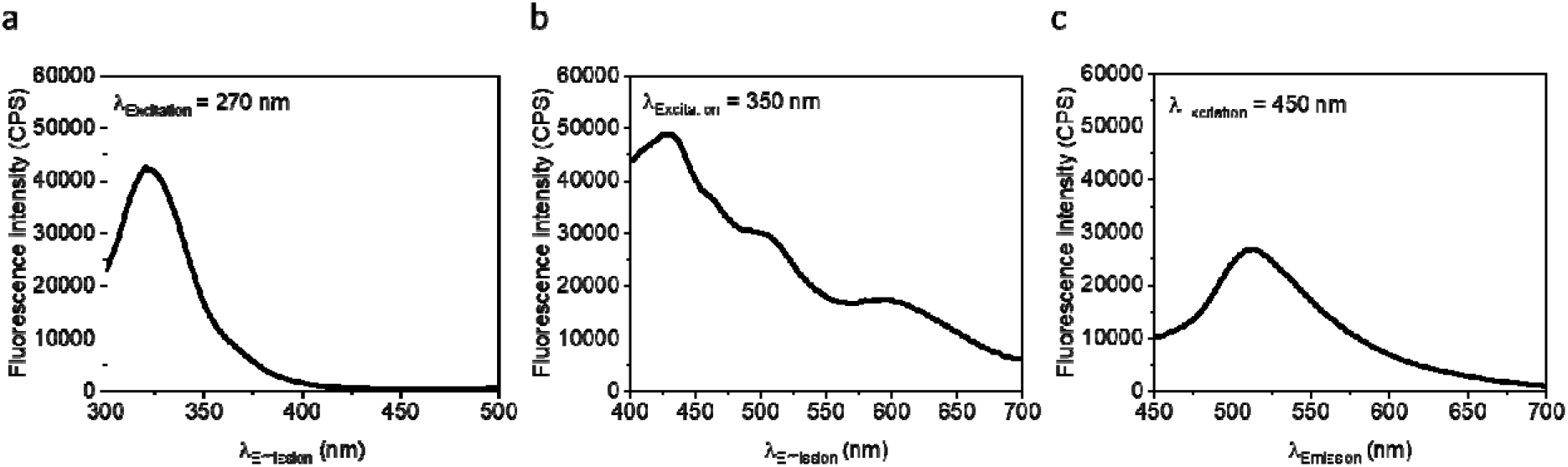
Spectral identification of fluorescent redox-active species. Fluorescence spectra of beet juice were measured at wavelengths compatible with proteins (consisting of tyrosine and tryptophan), NADH, and flavins. **a** Fluorescence spectra exciting at 270 nm which is compatible for protein detection. **b** Fluorescence spectra exciting at 350 nm which is compatible with NADH detection. **c** Fluorescence spectra exciting at 450 nm which is compatible with flavins detection.

### Dark and photocurrent generation from beet juice

Based on the analysis showing that beet juice consists of redox species, we next wished to assess whether the juice can be a source for bio-electricity generation in a BEC. Furthermore, previous studies showed that some bio-components such as benzoquinones or flavins[13] may also undergo photoelectrochemical reactions converting them into a reduced form that can transfer electrons to the anode to produce electrical current. To explore this, chronoamperometry (CA) measurements of beet juice were measured (Fig. 3). The measurements were conducted with dark/light intervals of 400 s with an applied potential of 0.9 V chosen based on the biggest peak obtained in the CV measurements (Fig. 1b). The obtained results showed a decaying current density pattern starting from ∼120 µA / cm^2^ down to ∼30 µA / cm^2^ after 400 s. upon white light illumination, a photocurrent of ∼3 µA / cm^2^ above the dark current was obtained. We suggest that some of this photocurrent originates from flavins and anthocyanins as previously reported for cherry juice[36]. CA of NaCl was measured as a control experiment showing no significant current generation in dark or light.

**Fig.3.**
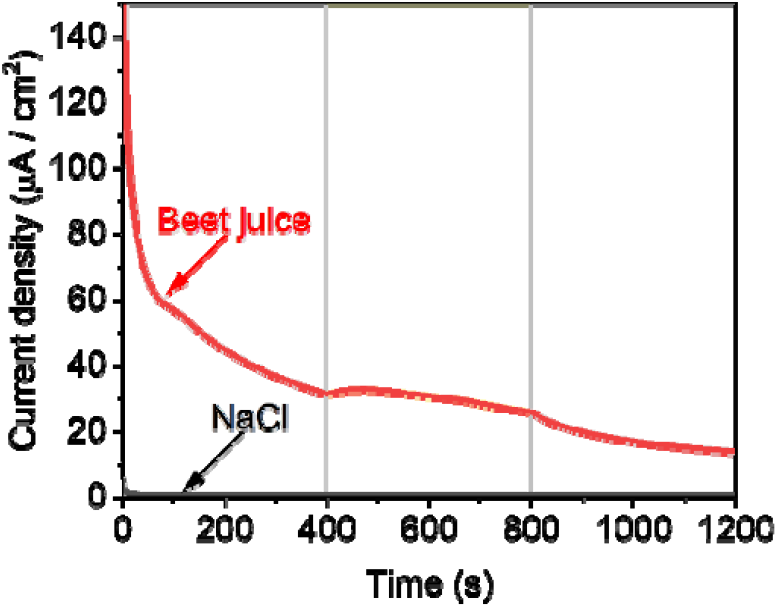
Dark and photocurrent generation from beet juice. CA of NaCl (black) and beet juice (red) were measured with an applied potential of 0.9 V with dark/light intervals of 400 s. Gray and light-yellow rectangular backgrounds represent dark and light respectively.

**Fig.4.**
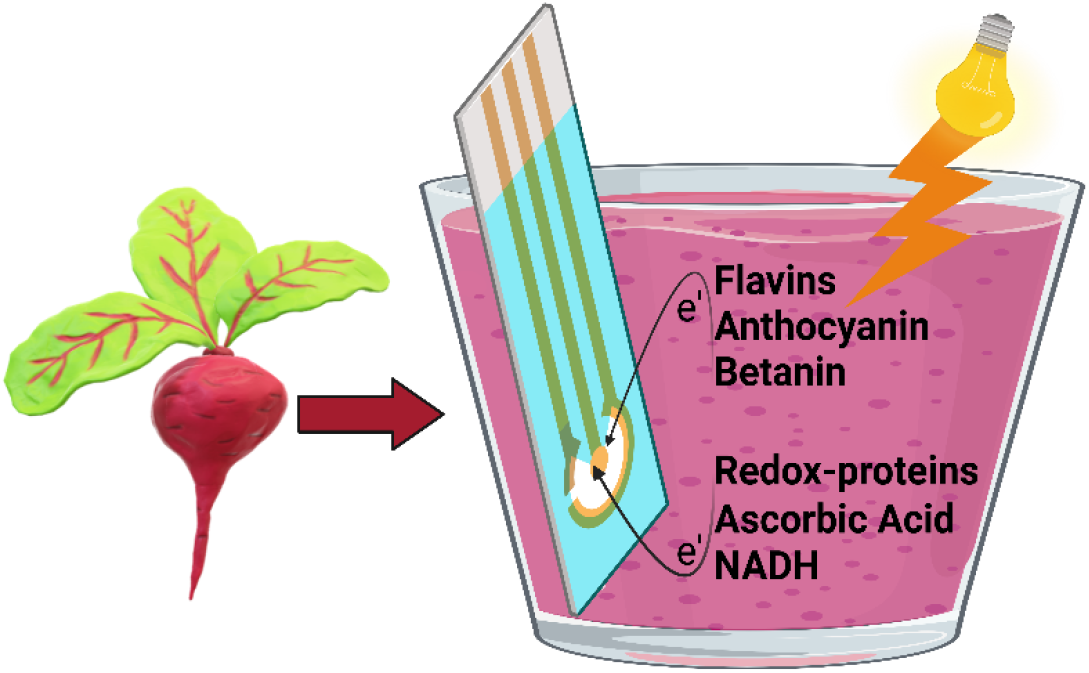
Proposing a mechanism for the dark and photocurrent generated by beet juice. The complex solution of beet juice consists of reducing molecules that can donate electrons at the anode of the BEC generating electricity. Among these species are redox-active proteins, ascorbic acid, and NADH. Upon white light irradiation, components such as flavins, anthocyanin, and betanin are photo reduced enabling them to become electron donors at the anode.

### Proposing mechanisms for the dark and photocurrent generated by beet

Based on the results obtained in this work, and previous studies of cherries-based BECs[36], we suggest a mechanism in which the major electron sources playing a role in the dark current production are NADH, and ascorbic acid, while part of the photocurrent may be generated by flavins[13] or pigments such as anthocyanin and betanin which were also reported to be active in dye-sensitized solar cells [45] Another potential contributor to the photocurrent may be reactive oxygen species (ROS)[46]. Nevertheless, we predict that the donation of ROS in beet juice is minor or even insignificant as their lifetime in the juice is expected to be very short.

## Conclusions

The development of methods for clean electricity production is highly important to fight future climate change disasters. In this work, we show the ability of beetroot juice to act as an electron source in a BEC. We apply electrochemical and fluorescence spectroscopy methods to identify the existence of NADH, ascorbic acid, flavins, and anthocyanin which are likely to play a major role in the dark and photocurrent production. Based on the observations of this work and previous work utilizing cherries in BECs to produce electricity, we suggest that the ability to use fruits and vegetables as an electron source is not specific and may be applied to many different species whose efficiency will be studied in the near future.

## Acknowledgments

We thank the University of California and a UCSB Faculty Research Grant for financial support. Yaniv Shlosberg is supported by the Otis Williams Fellowship. Fluorescence measurements in this study were obtained using central facilities at the UCSB Materials Research Laboratory (MRL), along with technical support from Jaya Nolt. Some of the figures were prepared using Biorender.com.

## Author Contributions

Y.S conceived the idea. Y.F, D.R and Y.S designed and performed the experiments. Y.F and Y.S wrote the manuscript.

